# MAL suppresses OSCC tumorigenesis by maintaining epithelial cell differentiation

**DOI:** 10.1101/2021.12.01.470749

**Authors:** Xueqin Zhu, Zheqi Liu, Shengcai Qi, Xin Zou, Tingwei Lu, Xing Ke, Xing Qin, Xiaoning Wang, Ming Yan, Qin Xu, Jianjun Zhang, Xu Wang, Zhen Zhang, Wei Cao, Xiangbing Wu, Wantao Chen

## Abstract

Oral squamous cell carcinoma (OSCC) is widely recognized as an optimal model for precise medicine guided molecular biomarkers of cancer, however, few clinical practices were applied till now. Based on the data from our own studies and published papers, it was found that the expression of *MAL* was significantly decreased in epithelial cancer as compared with normal tissues, and exhibited a opposite association with pathological grade. To study the molecular events related to deficiency of *MAL* during carcinogenesis, occurrence and development, a *Mal* knockout mouse model was constructed and consistently reproduced and bred. The *Mal* knockout mice are highly vulnerable to tumor induction by carcinogen of 4NQO, evidenced by their extremely earlier carcinogenesis, higher incidence, and more aggressive growth. Analysis of scRNA-seq data indicated that *Mal* knockout mice lost the ability in maintaining epithelial cell differentiation and get more prone to carcinogen with a remarkably higher incidence of epithelial malignancy. Further analyses identified putative co-functional genes of *MAL*, including *DSG1, AQP3* and *S100A8*, which are key factors in maintaining epithelial cell differentiation. To conclude, the current study exhibits the clinical significance and explains the tumor suppressing function of *MAL*. The results also suggest the potential of *MAL* and its co-functional genes being biomarkers for designing the prevention and/or differentiation therapy strategies in OSCC.

**Significance:** *MAL* is found to be strongly opposite with tumor pathological grade from clinical and in vivo studies in OSCC. We propose *MAL* and its co-functional genes, including *DSG1, AQP3* and *S100A8*, as key factors in maintaining epithelial cell differentiation and are valuable targets for designing prevention and differentiation therapy strategies in OSCC.

## Introduction

Oral squamous cell carcinoma (OSCC) is the most common cancer of head and neck region obtained from mucosa epithelial cells, which accounts for more than 90% of all oral cancers, with the main risk factors being the consumption of tobacco and/or alcohol and chewing areca^1,2^. Despite ongoing efforts, few effective biomarkers and targeted therapeutic strategies have been identified or developed yet for prevention or treatment in the patients with OSCC. Therefore, a fully comprehensive understanding of the molecular pathogenesis of OSCC are crucial to the interceptive attempts against this kind of malignancy.

To further explore the key molecules in the carcinogenesis, occurrence, and progression in OSCCs, several candidate genes were filtered according to microarray of OSCC and paired normal epithelial tissues^3^. Among these genes, we preliminarily investigated the functions of myelin and lymphocyte protein (*MAL*) in OSCC and found *MAL* played a role of tumor suppressor in OSCC^4^. *MAL* is located on human chromosome 2, 2q13 and includes 4 exons and 3 introns. A study found that the protein encoded by *MAL* works as a proteolipid protein which is involved in apical transport machinery and signal transduction of epithelial cells^5^. Recently it has been reported as a tumor suppressor in several kinds of epithelial cancers^6-12^. Our previous studies suggested that *MAL* expression had a close relationship with the pathological grade of OSCC. However, evidence of how *MAL* can suppress the epithelial cancer and affect the pathological grade or the mechanisms behind this, is still so lacking.

There is no doubt that the establishment of clear new animal model of OSCC, which has high similarity with human in phenotypes, especially molecular genetic phenotypes, will help to develop a novel mechanism, diagnosis, prevention and treatment of OSCC. To address these questions, a *Mal* knockout mouse model was established and 4-nitroquinoline-1-oxide (4NQO) was applied to induce tumors in the tongue, which could mimic the interactions among molecular events and environment factors in human OSCC development. Single-cell transcriptome was used to explore the specific molecular mechanism of how *MAL* affecting the carcinogenesis, occurrence and progression of OSCCs.

## Results

### *MAL* expression level oppositely correlated with OSCC pathological grade

In our previous studies, we screened out a serious of genes that related to the progression of OSCC through oligonucleotide microarray (Affymetrix Hk-4U 95AV2). Among them, the decreased expression of *MAL* in tumors caught our attention (Figure 1a). Through the analysis of pan-cancer data from TCGA, we found that *MAL* expression significantly down-regulated compared with normal tissues in most cancer types (Figure 1b). Next, we analyzed *MAL* protein expression in different human tissues through The Human Protein Atlas (HPA https://www.proteinatlas.org/). The results indicated that in head and neck, *MAL* was highly expressed in esophagus and tongue tissues (Figure 1c). Furthermore, we performed RNA-seq using 100 tumor tissues and 99 adjacent normal mucosae of OSCC patients from the Shanghai Ninth People’s Hospital, Shanghai Jiao Tong University School of Medicine. The receiver operating characteristic (ROC) analysis revealed that *MAL* transcription expression could be used as a biomarker to distinguish tumor from non-tumor in Shanghai Ninth People’s Hospital cohort (9H-cohort) and TCGA-cohort with AUC of 0.950 and 0.866, respectively (Figure S1a). Clinical correlation analysis showed that *MAL* transcription expression was oppositely correlated with tumor pathological grade (Figure 1d), which was also verified in TCGA-cohort (Figure 1e), while the correlation between *MAL* transcription expression with T or N stage was not consistent in two cohorts (Figure S1 b and c). The results of immunohistological chemistry (IHC) also showed the protein expression level of *MAL* in normal mucosae was much higher than that in tumor tissues, and the expression in pathological grade I tumors was higher than those in grade II and grade III tumors (Figure 1f). These results suggested that *MAL* is closely related to epithelial cell differentiation in OSCCs.

**Figure 1.**
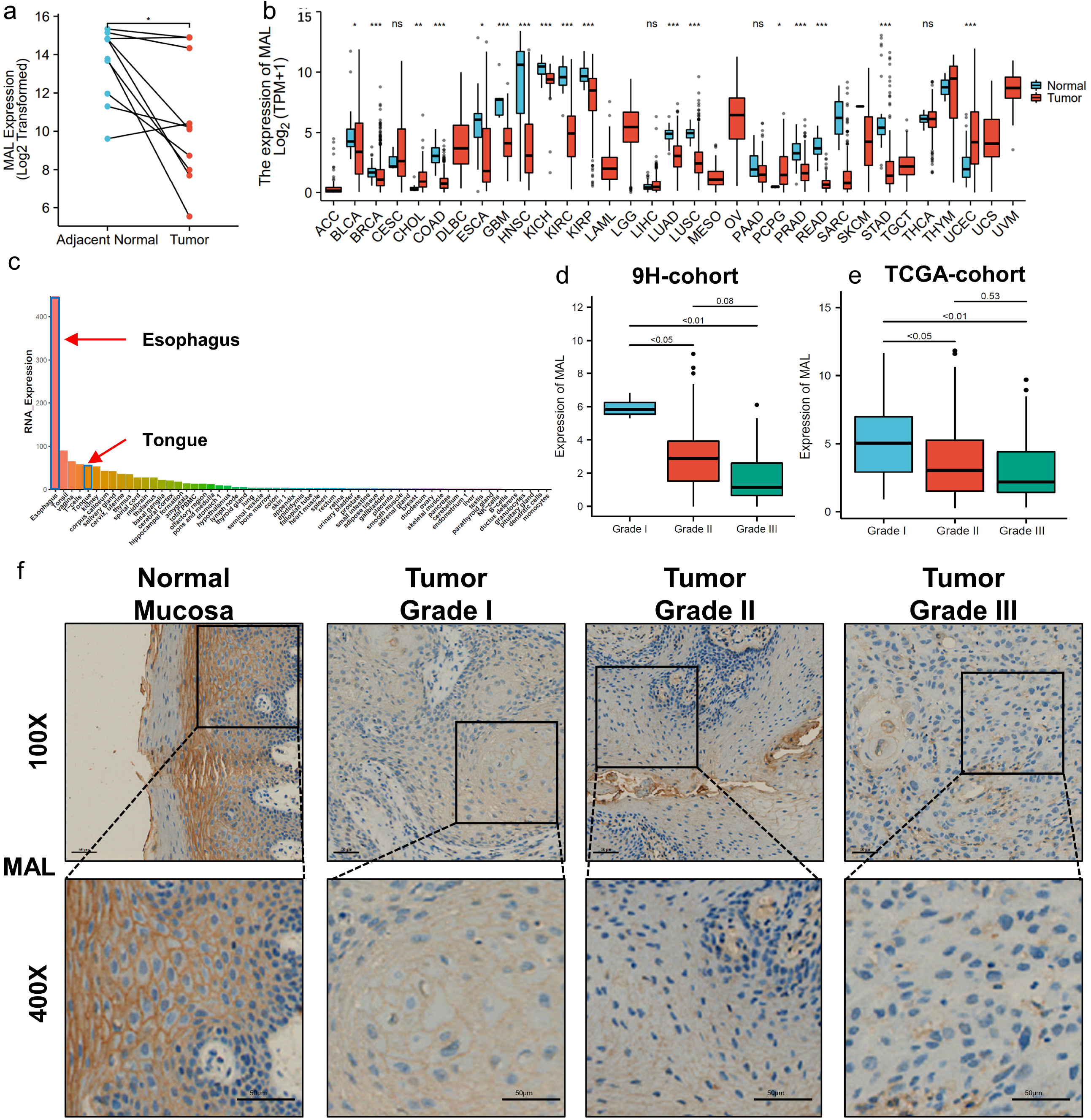
*MAL* expression is oppositely related with tumor pathological grade in OSCC. a. The *MAL* transcription expression in 10 paired normal and tumor tissues using microarray. b. The *MAL* transcription expression in pan cancer from TCGA. In most cancer types, *MAL* expression in tumor tissues is lower than in normal tissues. c. *MAL* protein expression is enriched in esophagus and tongue mucosa from HPA database. d. The association between tumor pathological grade and *MAL* transcription expression from 9H-cohort. e. The association between tumor pathological grade and *MAL* transcription expression from TCGA-cohort. f. IHC slides show the protein expression of *MAL* is oppositely correlated with tumor pathological grade from 9H-cohort. \**P* < 0.05, ** *P* < 0.01, *** *P* < 0.001

### Establishment and phenotype identification of *Mal* knockout mice

The *Mal* knockout mice were successfully constructed and mice that had the *Mal* knocked out were still able to produce offspring. Before the tumor chemical induction process, no significant difference was observed in appearance, weight and hair between *Mal* knockout and wildtype mice. The *Mal*-/- and *Mal*+/+ mice showed no spontaneous tumor in organ or tissue other than oral mucosa (Figure 2c and e). HE staining was performed on the oral mucosa, and no obvious abnormality was observed in the microstructure of the tongue mucosa of mice after *Mal* knockout, so as the proliferation related protein Ki67 and cell cycle related protein cyclin D1 (Figure 2d). Meanwhile, no significant difference was seen in blood routine or chemistry between normal breeding *Mal*-/- and *Mal*+/+ mice (Figure S2b and c). Through the examination of the tissue sections of the mouse esophagus, spleen, thymus, kidney, liver, lung, skin, stomach and testis, no obvious structural abnormalities were found (Figure 2f). The levels of AST, ALT, ALB, TP, GLO, A/G, BUN, CREA and GLU in mice blood was also detected, indicating that *Mal* knock out makes no significant difference under physical conditions (Figure S2a). In addition, result from flow cytometry showed that the proportion of T cells, B cells, and NK cells from peripheral blood and spleen had no significant difference between *Mal*-/- and *Mal*+/+ mice, indicating a normal immune system of *Mal-/-*mice (Figure S3 and S4). These results suggested that knockout of *Mal* did not make a significant impact on normal physical indexes in mice.

**Figure 2.**
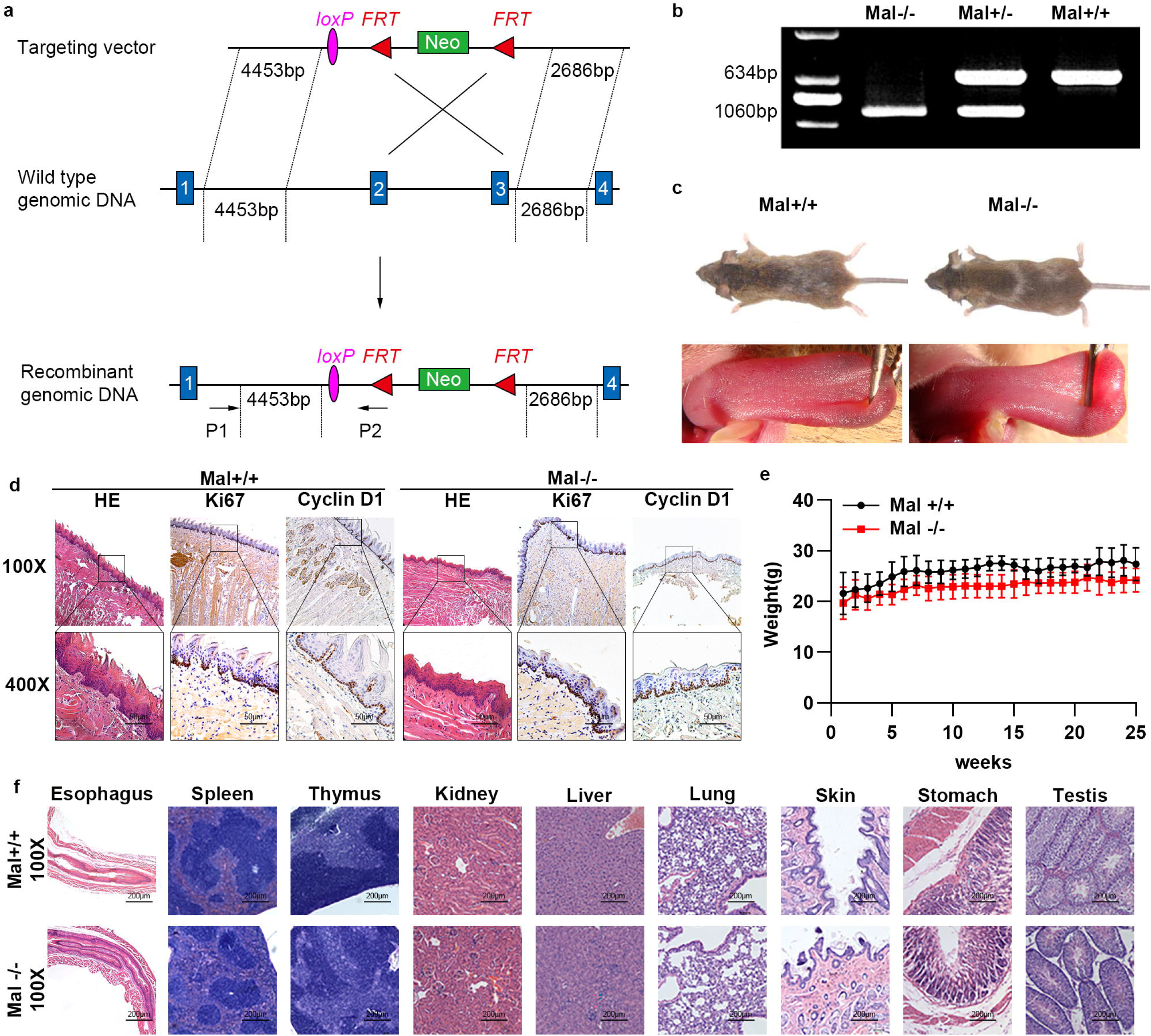
*Mal* knock out mice were successfully constructed. a. The design strategy of targeting vector to knock out 2 and 3 exons of *Mal* gene in mouse. b. Agarose gel electrophoresis shows the genotype of *Mal-/-, Mal+/-* and *Mal+/+* mice. c. No obvious changes, such as body size, hair and tongue mucosa, are observed after *Mal* knockout. d. The HE shows that no obvious abnormality is found in the microstructure of oral mucosa in *Mal*-/-mice compared with *Mal*+/+ mice, so as the proliferative markers Ki67 and cyclin D1. e. The change trend of body weight of *Mal*-/- and *Mal*+/+ mice under normal physiological conditions. f. *Mal* knockout does not affect the microstructure of esophagus, thymus, spleen, kidney, liver, lung, skin, stomach and testis.

### *Mal* depletion promoted aggressiveness of OSCC induced by 4NQO

The *Mal* knock out and wild type mice were fed with drinking water containing 4NQO at a dose of 40 or 80 µg/mL, the changes of oral mucosa and gloss of each mouse were strictly observed and the exact occurrence time of oral mucosal tumors was recorded once a week. When induced by a lower dose of 40 µg/mL, the first oral lesion was observed in *Mal-/-*mouse on tongue mucosa at 10^th^ week, while the first oral lesion in *Mal*+/+ mouse occurred 2 weeks later. At week 22, only 27% (3/11) *Mal+/+* mice developed oral lesions but the occurrence was up to 75% (6/8) in *Mal-/-*mice (Figure 3b and 3c). At the same time, the number of oral tumors in mice were counted and there were more oral tumors in *Mal-/-*mice than *Mal+/+* mice (*P=0*.*0413*) (Figure 3c). The results of HE showed that most of the oral lesions in *Mal+/+* mice were pre-cancer or carcinoma in situ, which did not break through the epithelial basement membrane, with obvious keratinization and a small amount of mitosis under a lower induction dose of 40 µg/mL. However, after *Mal* knockout, the oral tumors had obvious exophytic growth, and most of them were obviously infiltrating growth, which broke through the epithelial basement membrane, with high atypia and small nuclei. The IHC staining of cell proliferation related protein Ki67 and cell cycle related protein cyclin D1 showed a higher IHC score in *Mal-/-*mice than *Mal+/+* mice (Figure 3d-f). Similarly, when the 4NQO-induced dose was increased to 80 µg/mL, the time of oral mucosal lesions occurred in the *Mal* knockout mice were significantly earlier, and the number of oral tumors was much more than that in WT mice (Figure 4a-d). HE and IHC observations also showed that oral tumors of *Mal* knockout mice were more aggressive and had lower epithelial differentiation (Figure 4e and S5). To sum up, *Mal* knockout can elevate the susceptibility of oral mucosal epithelium to carcinogen, shorten the carcinogenesis period and increase oral tumor incidence, which reveal a more intensive invasion and lower differentiation level for low induction doses (40 and 80µg/mL).

**Figure 3.**
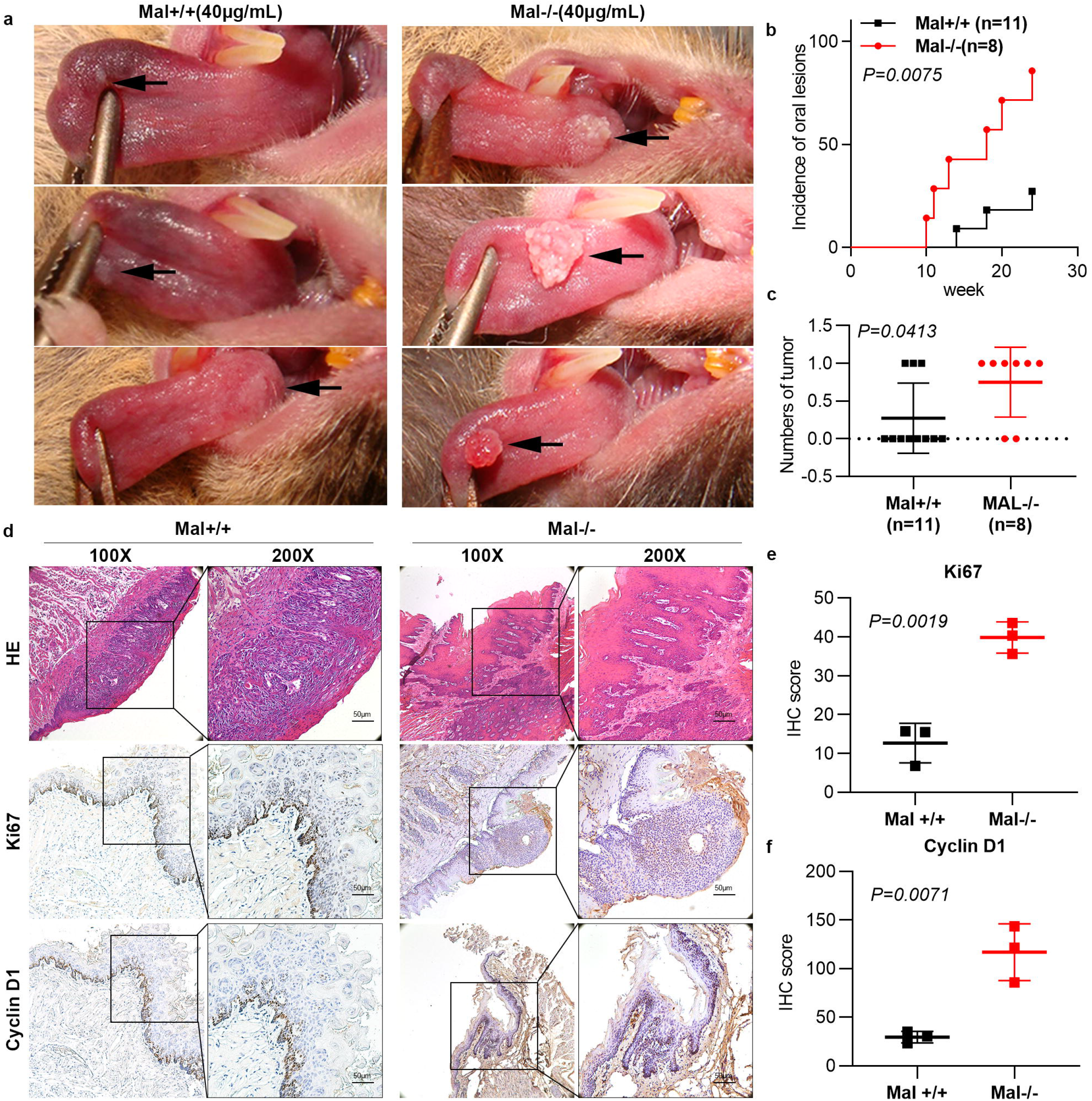
Phenotype of oral mucosa induced by 4NQO at a dose of 40 μg/mL in *Mal* knockout mice. a. At the 24^th^ week of observation, the oral mucosa of *Mal+/+* mice shows white plaques, while in *Mal-/-*mice, the oral mucosal tumors of mice show obvious exogenous growth and cauliflower shape. b. The incidence of oral mucosal tumors in *Mal-/-*mice is significantly higher than *Mal+/+* mice, and the occurrence time is earlier (*P=0*.*0075*). c. The number of oral tumor occurred in *Mal-/-*mice is much more than *Mal+/+* mice (*P=0*.*0413*). d. The histopathological feature of oral mucosal tumors from *Mal-/-* and *Mal+/+* mice. e and f. *Mal-/-*mice have higher IHC score of Ki67 and cyclin D1 than *Mal+/+* mice.

**Figure 4.**
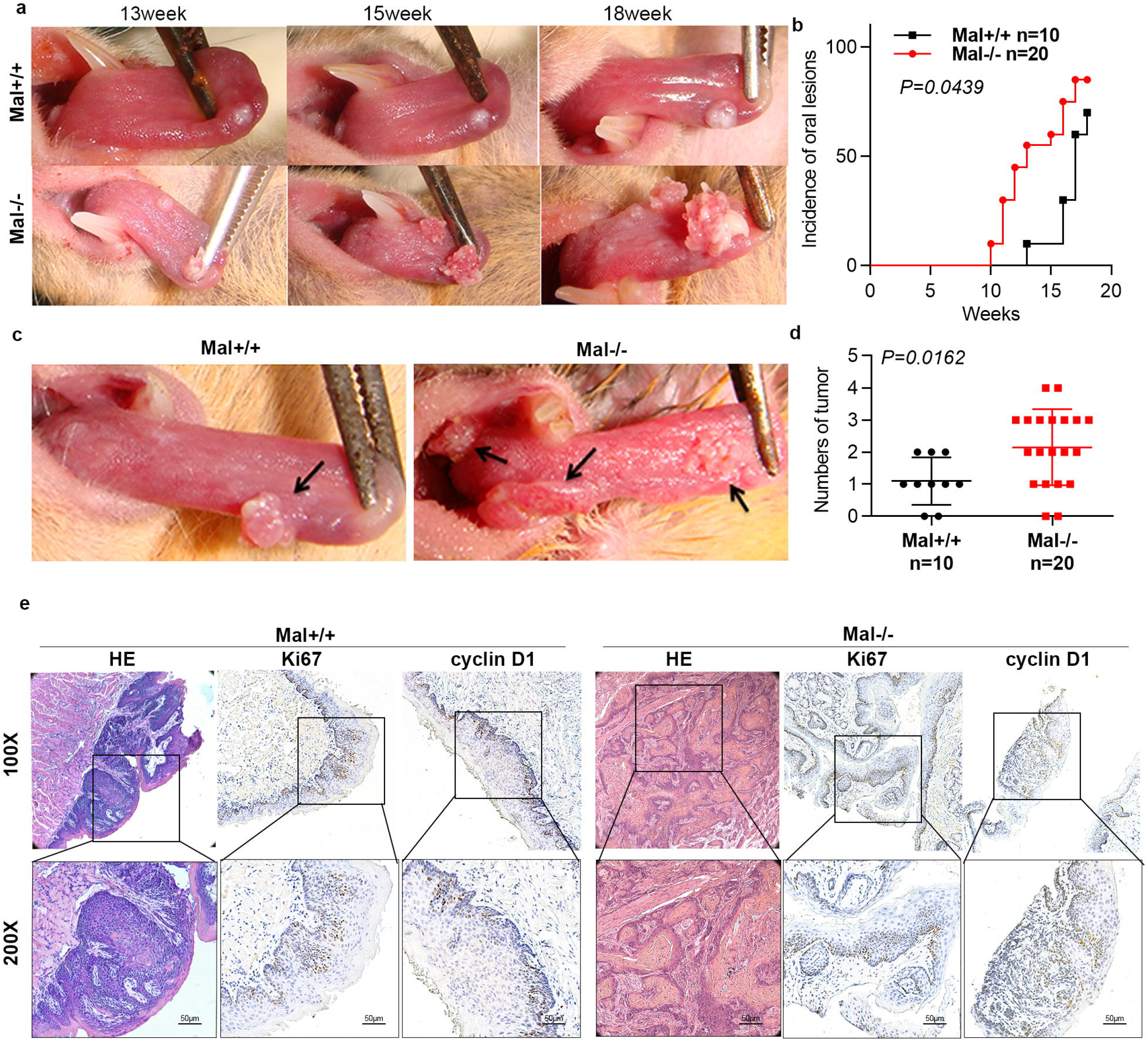
Phenotype of oral mucosa induced by 4NQO at a dose of 80 μg/mL in *Mal* knockout mice. a. The oral tumors of *Mal+/+* and *Mal-/-*mice are observed and photographed at the 13^th^, 15^th^ and 18^th^ week, respectively, and it is found that the oral tumors of *Mal-/-*mice are larger and grow faster than *Mal+/+* mice at the same time point. b. The incidence of oral mucosal tumors in *Mal-/-*mice is significantly higher than *Mal+/+* mice, and the occurrence time is earlier (*P=0*.*0439*). c and d. In most *Mal+/+* mice, only one oral tumor is seen, while more than two oral tumors are found in 70% (14/20) *Mal-/-*mice. The number of oral tumors occurred in each *Mal-/-*mice is much more than *Mal+/+* mice (*P=0*.*0162*). e. The histopathological feature of oral mucosal tumors from *Mal-/-* and *Mal+/+* mice.

### scRNA-seq indicated *Mal* suppresses tumorigenesis via maintaining epithelial differentiation capability

Seven tumor samples, including 3 tumors from WT mice and 4 from *Mal* knockout mice were collected. scRNA-seq was conducted for these samples via the 10× Genomics platform. Unsupervised clustering analysis of 74940 cells from 7 tumor samples revealed 28 distinct clusters (Figure 5a left). According to the expression pattern of canonical markers, cell types were annotated (Figure 5a right, Figure S6a). It was shown that *Mal* was mainly expressed in epithelial cells (Figure 5b), thus the epithelial cells were interrogated in the subsequent analysis. The epithelial cells were further categorized into nine clusters with distinctive gene expression profiles (Figure 5c and 5f), among which, cluster 5 was almost consisted of cells from WT mice (Figure 5c and 5d). Single cell CNV analysis illustrated all epithelial cell clusters consisted of both malignant and non-malignant cells, but with different preferences (Figure S6b-e). In addition, the expression of *Mal* was enriched in cluster 5 and 7 compared to other clusters (Figure 5e). Since the cell number of cluster 7 was inadequate to represent the characteristics, cluster 5 fit for further investigation. To speculate the relationship between cluster 5 and tumorigenesis, the top 30 gene markers (with largest fold changes) of cluster 5 were used as a gene panel. On TCGA-cohort^13^, Kaplan-Meier survival analysis indicated that high expression of this gene panel significantly associated with a better survival rate (Figure 5g). The trend was consistent with the function of *Mal* in inhibiting carcinogenesis and development, which was validated in the clinical samples.

**Figure 5.**
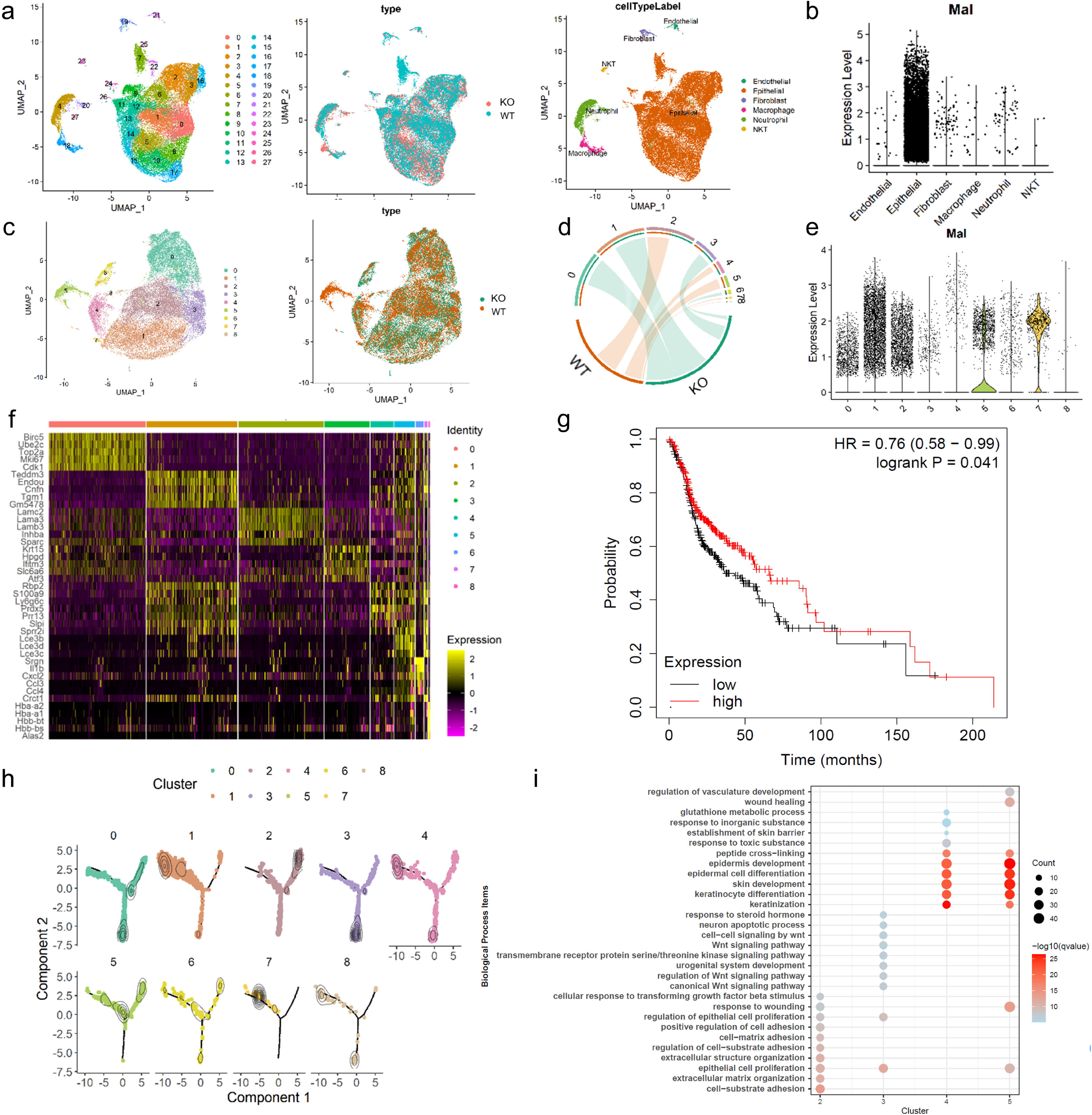
*Mal* suppresses tumorigenesis via maintaining epithelial differentiation via scRNA-seq. a. (a) Uniform manifold approximation and projection (UMAP) of scRNA-seq cells recovered from tumor tissues of mice. From left to right: cells are labeled by unsupervised clustering, type of mice and cell types. b. The violin plots of the *Mal* expression in different cell types, which show a relatively high expression in epithelial cells. c. The UMAP of epithelial cells, left is labelled by unsupervised clustering and the right is labelled by type of mice. d. Chord diagram shows the different cell clusters of epithelial cells are belonged to different mice. e. The violin plots of the *Mal* expression in different cell types of epithelial cells, which show a relatively high expression in cluster 5 and cluster 7. f. The heatmap shows the distinction and marker genes of different clusters in epithelial cells. g. Using the top 30 gene markers of cluster 5 as a gene set, the Kaplan-Meier plot shows the high expression of cluster 5 gene set has a better survival in TCGA. h. Three trajectory end nodes are distinguished using pseudotime trajectory analysis. The cluster 5 is localized in the intersection of three nodes. The representative cell cluster of three nodes are cluster 2 (the upper right one), cluster 4 (the upper left one) and cluster 3 (the lower one), respectively. i. The bubble plot shows biological process of cluster 2, 3, 4 and 5. The cluster 5 has a similar function with cluster 4 which is enriched in epithelial differentiation.

To further study the characteristics of epithelial cells, pseudotime trajectory analysis^14^ was performed. It was observed that there were three trajectory end nodes, each of which was dominated by cluster 2, 3 and 4, respectively (Figure 5h). Furthermore, cluster 5 located in the middle position of the trajectory but closer to cluster 4. As it has been shown in Figure 4d, cluster 2 and 4 were dominated by cells from WT mice and cluster 3 from *Mal* knockout mice, these three end nodes may represent distinctive characteristics of tumor cells in WT and *Mal* knockout mice, respectively. Such speculation was further supported by the GO analysis on the markers of each of the 3 clusters (Figure 5i). It was found that cell adhesion and epithelial differentiation were mostly enriched in cluster 2 and cluster 4, which has been known to counteract tumorigenesis in OSCC. On the contrary, Wnt signaling pathways were enriched in cluster 3, which were the classic oncogenic factors in OSCC. Such observation is consistent to the results presented in the previous sections, which implied deficiency of *Mal* may promote OSCC tumorigenesis. To be noticed, the function of cluster 5 was more similar with that of cluster 4 which was related with epithelial differentiation (Figure 5i). Considering cluster 5 was mainly from WT mice, the results suggested that deficiency of *Mal* may promote tumorigenesis by preventing epithelial cell differentiation.

### Co-functional genes of *Mal* in regulating epithelial differentiation

To further investigate the potential mechanism from which *Mal* regulates epithelial differentiation, we managed to identify key genes that were closely related with *Mal*. Firstly we selected the top ten functional gene sets of cluster 5. The Homo Sapiens homologous genes of selected gene sets were interrogated on a TCGA-cohort. Eight out of 10 gene sets were found to be positively correlated with overall survival (Figure 6a and S7a). By overlapping the marker genes included in the eight gene sets, we finally got three genes, i.e., *ERRFI1, DSG1* and *AQP3* (Figure 6b). In addition, we also investigated the top 10 Homo Sapiens homologous genes cell markers of cluster 5 cells, i.e., *SLPI, S100A8, LCE3B, LCE3D, LCE3C, S100A9, HIST1H1B, MARCKSL1*, and *CXCL12*. Using TCGA-cohort, we calculated the Spearman correlation between *MAL* and each of those genes, identified *DSG1, AQP3, SLPI, S100A8, LCE3B, LCE3C, LCE3D* and *S100A9* (Figure 6c). Further validation using mouse scRNA-seq data, TCGA-cohort, and 9H-cohort RNA-seq data, *DSG1, AQP3* and *S100A8*, having all positive correlation with *MAL* in three datasets, were finally screened out as the co-functional genes (Figure 6d and S7b). The relationship between 3 co-functional genes and tumor pathological grade was further demonstrated on the RNA-seq data from TCGA-cohort and 9H-cohort, i.e., with the increase of tumor grade, the expression of *DSG1, AQP3* and *S100A8* decreased gradually (Figure 6e). Such relationship was also verified by IHC observation (Figure 6f). In summary, evidences from previous researches and our study hereby demonstrated *MAL* and co-functional genes play a key role in maintaining differentiation capability of epithelial cells.

**Figure 6.**
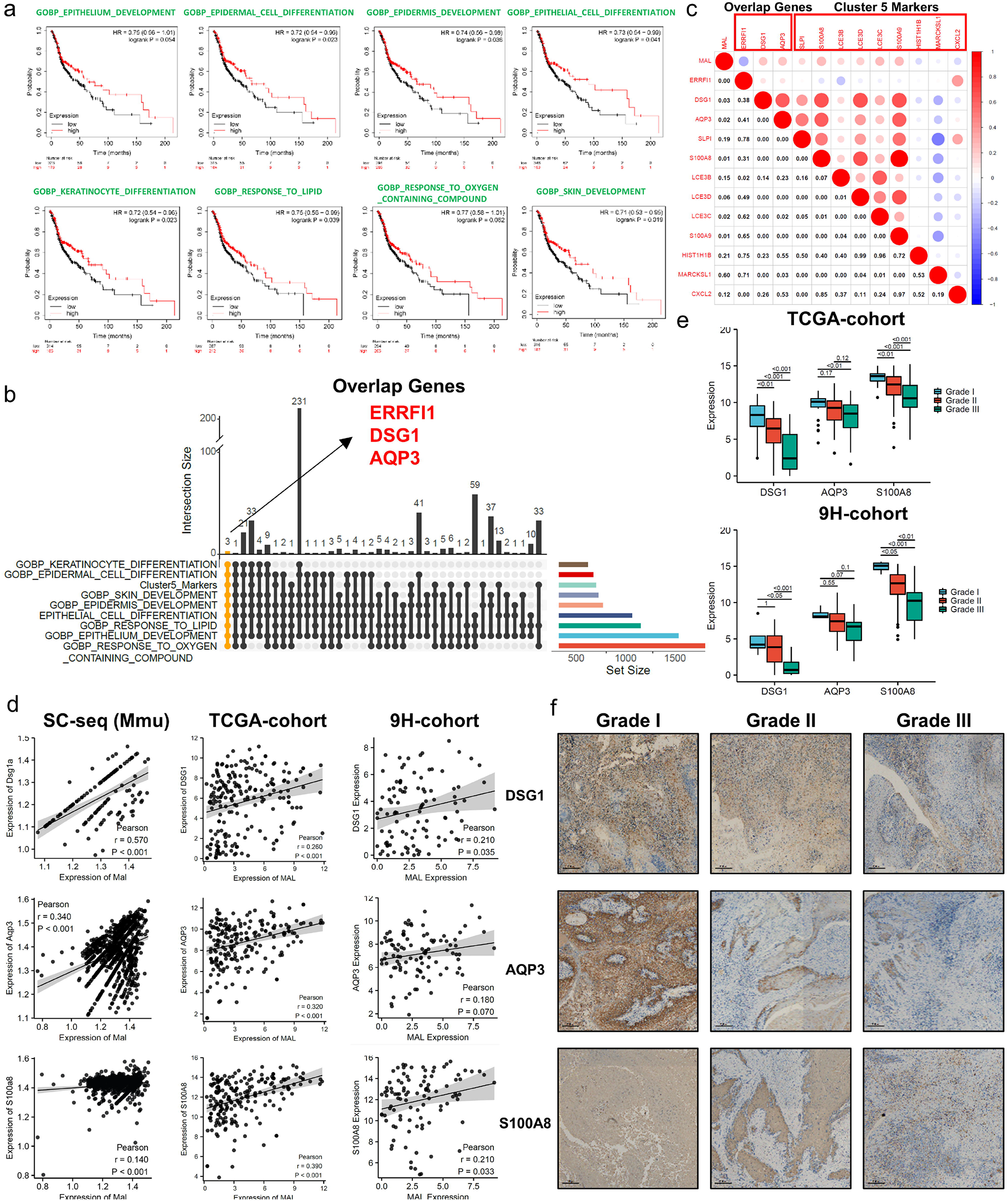
*DSG1, AQP3* and *S100A8* serve as co-functional genes with *MAL* in regulating epithelial differentiation. a. The genes derived from 8 out of 10 functional pathways are performed by Kaplan-Meier analysis, which predict a better survival in TCGA-cohort. b. Upset plot shows the process of screening out candidate genes. Three candidate genes, i.e., *ERRFI1, DSG1* and *AQP3*, are picked out as shown. c. Combined the candidate genes from upset plot and 10 gene markers of cluster 5, the correlation heatmap shows *MAL* has a positive correlation between *DSG1, AQP3, SLPI, S100A8, LE3B*-*D* and *S100A9*. The Spearman correlation analysis is performed using TCGA-cohort. d. Dot plots show the correlation between *MAL* and *DSG1, AQP3* and *S100A8* in scRNA-seq data, TCGA-cohort and 9H-cohort. These genes are all positively correlated with *MAL* in three datasets and are identified as co-functional genes. e. The association between tumor pathological grade with *DSG1, AQP3* and *S100A8* transcription expression from TCGA-cohort and 9H-cohort. As the pathological grade rise, the expression of three genes is decreased. f. IHC slides show the protein expression of *DSG1, AQP3* and *S100A8* are oppositely correlated with tumor pathological grade from 9H-cohort.

## Discussion

Recent studies showed that *MAL* tended to tone down in many epithelial malignancies, such as esophageal cancer^6^, cervical cancer^7^, colon tumor^8^, gastric cancer^9^, breast cancer^10^, salivary gland cancer^11^ and non-small cell lung cancer^12^. *MAL* was mostly considered as tumor suppressor gene in epithelial tumor. Our previous study revealed that *MAL* was significantly down-regulated in OSCC and correlated with tumor growth^4^. Further in this study, we analyzed the relationship between *MAL* expression and tumor pathological grade using tumor samples from our institution and TCGA data. The results of both data confirmed that *MAL* transcription and protein expression were oppositely correlated with the pathological grade of OSCC, which shed light on further investigation of *MAL* function in tumorigenesis.

OSCC originates from the abnormal differentiation and proliferation of oral mucosal epithelium. The development of OSCC exhibit consistent invasion and infiltration to peripheral normal tissues, which is characterized by a gradually increased aggressiveness along the process. Useful and effective animal models are optimal tools for better understanding such long-term and multistep progression. The successful construction of 4NQO-induced OSCC model takes at least 6 months, which can mimic the epithelial malignant transformation progress in human OSCC. With a serious of in silico, in vitro and in vivo attempts, the genetic modified mouse model exactly described the vital role of *Mal* in guaranteeing the epithelial cell to classic cell fate. In our study, the malignant features were markedly enhanced as validated by the differences regarding to early tumor onset, obvious infiltrating growth, more number of lesions, as well as high staining intensity of proliferative markers. These findings again emphasized the importance of regular epithelial differentiation sustained by *Mal*.

To further explore the underlying mechanism, scRNA-seq was performed. The results of scRNA-seq data also confirmed the close relationship between *Mal* expression and epithelial cell differentiation, which was consistent with our previous clinical analysis and in vivo results. According to the comprehensive analysis and cross validation based on scRNA-seq of mouse tumors, RNA-seq data from our institution and TCGA, we found out and proved 3 co-functional genes with high correlation with *MAL*.

In humans, *S100A8* and *S100A9* typically form heterodimers (*S100A8*/*A9*), which is called calprotectin^15^. This complex appears to be a tumor-suppressor and reduction in its protein expression is associated with poor prognosis in HNSCC^16-18^. What’s more, a study indicated that *S100A8*/*A9* is directly associated with cellular differentiation and appears to promote caspase-3/7-mediated cleavage of EGFR in HNSCC^19^. Aquaporin-3, encoded by *AQP3*, is the most abundant aquaporin in human epidermis^20^. While in the studies of oral mucosa, it was found that with the progress of oral mucosa epithelium to hyperplasia or even malignant transformation, the expression of *AQP3* gradually decreased^21^. Desmogleins (DSGs) work as desmosome components, are critical for the structure of intercellular junctions and maintenance of epithelial barrier integrity. They are also the key regulators of cell differentiation, epidermal homeostasis and carcinogenesis. Loss of desmosome proteins will promote the process of epithelial-mesenchymal transition (EMT)^22^. As a member of this family, the decreased expression of *DSG1* has been reported to be closely related to the occurrence of tumors and poor prognosis in a variety of tumors, including lung cancer, esophageal squamous cell carcinoma, extrahepatic cholangiocarcinoma, anal carcinoma, etc^22-25^. The studies in OSCC also suggested that the abnormal decrease of *DSG1* leads to the loss of intercellular connectivity, thus promoting tumor progression^26,27^. In our study, the correlation between the co-functional genes and tumor pathological grade was validated by scRNA-seq data, TCGA-cohort and 9H-cohort, i.e., the tumor pathological grade was oppositely correlated with both transcription and protein expression of 3 co-functional genes. The evidences from published studies and our results proposed *MAL* and its co-functional genes play a key role in maintaining differentiation capability of epithelial cells.

The results of this study also suggested that the decrease in *MAL* expression indicates the deterioration of epithelial differentiation, and tumor is more likely to occur, which is of remarkable for clinical prevention strategies, especially for patients with oral pre-malignant lesions. The primary goal of oral pre-malignant lesions management includes prevention, early detection, and treatment before malignant transformation^28^. The abnormal down-expression of *MAL* in oral pre-malignant lesions may indicate that the lesions are more likely to transit into OSCC, which reminds clinicians to deal with the lesions as early as possible.

Changes of *MAL* expression in tumor tissues could also provide new insights for tumor treatment in OSCC. Differentiation therapy is an effective treatment in some malignancies. The concept of differentiation therapy emerged from the fact that hormones or cytokines may promote differentiation ex vivo, thereby irreversibly changing the phenotype of cancer cells^29^. In hematologic malignancies, especially in acute promyelocytic leukemia, differentiation therapy is widely used^29,30^, but in solid tumors, the effect of differentiation therapy is still unsatisfactory^31^. The deficiency of *MAL* in OSCCs represents poor differentiation of cancer cells, which suggested we might reverse the process of tumor dedifferentiation by rebooting regulatory network of *MAL*.

In conclusion, we initially identified the role of *MAL* in the occurrence of OSCC and proposed potential regulatory factors of maintaining epithelial cell differentiation through which *MAL* affects tumorigenesis. The results also suggested the clinical application of *MAL* as a candidate target in designing treatment strategies. Further clinical studies are required to verify the clinical value of prevention or differentiation therapy strategies by targeting *MAL* and its co-functional genes (Figure 7).

**Figure 7.**
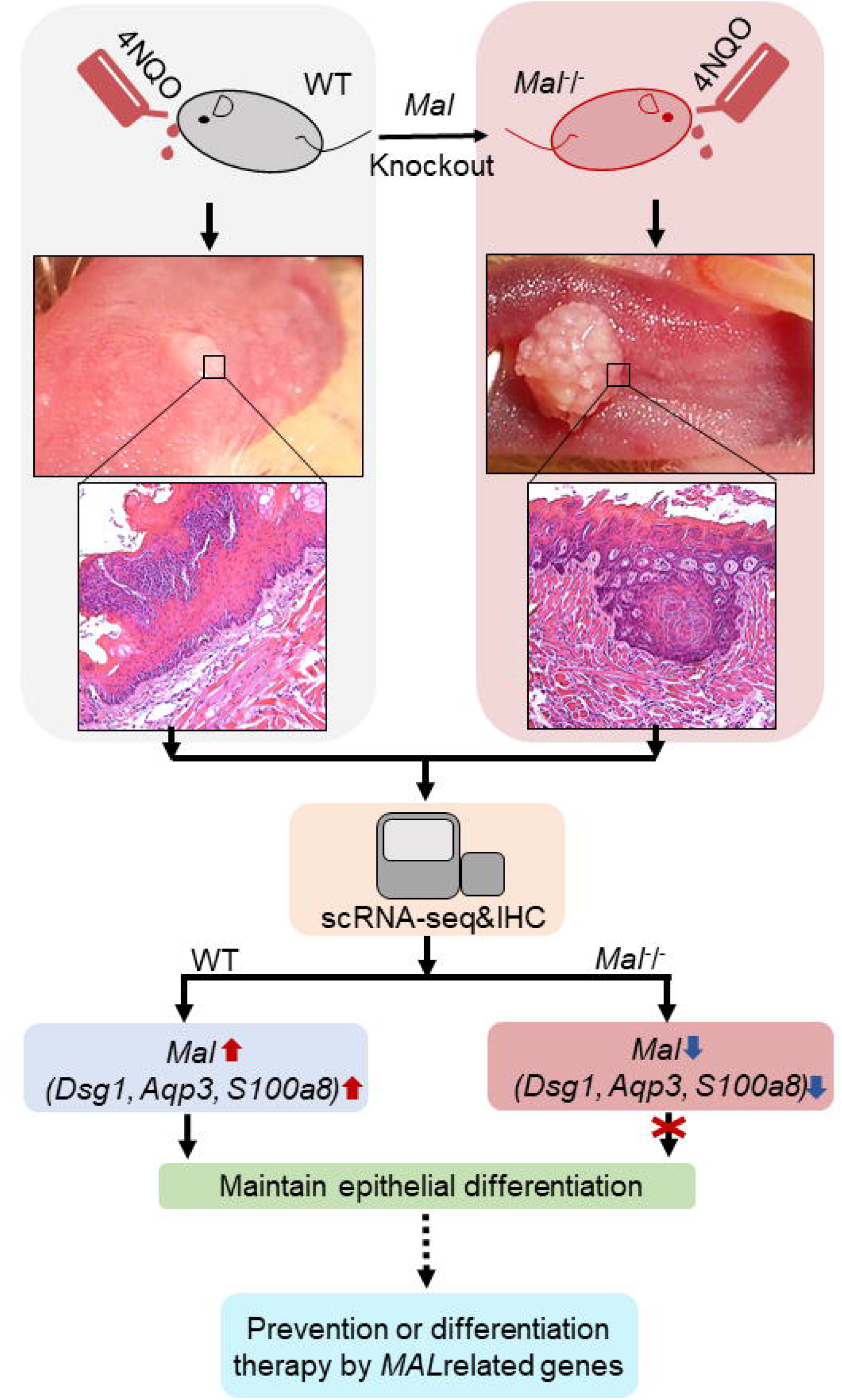
Schematic diagram showing genetic modified model and scRNA-seq identified *MAL* as a tumor suppressor of OSCC through maintaining epithelial differentiation.

## Materials and Methods

### Data acquisition and processing

Gene expression data, including the count and fragments per kilobase of transcript per million mapped reads (FPKM) and related clinical information of patients from TCGA HNSC project was downloaded from the UCSC Xena online database (http://xena.ucsc.edu/). FPKM data was transformed into transcripts per million reads, for further analyses. Patients with tumor in tongue, base of tongue and floor of mouth were picked up for further analysis and a total of 207 cases were finally selected. The microarray data (Affymetrix Hk-4U 95AV2) of 10 paired tumor tissues was collected from Shanghai Professional Technical Service Platform of Oral and Maxillofacial Tumor Tissue Samples and Bioinformatics Database.

### Sample collection

The tissue samples for RNA sequencing and paraffin-embedded samples of OSCC were collected from patients at the Ninth People’s Hospital, Shanghai Jiao Tong University School of Medicine. All patients involved were informed about the study and provided written informed consent, and the process was approved by the Ethics Committee of the Ninth People’s Hospital, Shanghai Jiao Tong University School of Medicine.

### Animal procedures

The *Mal* knockout mice were constructed in Shanghai Research Center of the Southern Model Organisms. All animal procedures were subjected to institutional ethical review and performed under the terms of Shanghai Jiao Tong University. The two transcripts of *Mal* was found from the Ensemble website (http://www.ensembl.org/Mus_musculus/Gene/Summary?g=EnsmusG00000027375). *Mal*+/-mice at the age of 8 weeks were caged to produce different *Mal* genotypes of offspring mice, providing sufficient numbers of mice for this study. The targeting vector was designed in the second and third exon according to the number and location of the exons of *Mal*. According to the design strategy, Neomycin (Neo) gene, a positive selective marker gene, was used to replace two exons of *Mal* sequence, and certain gene sequences to be deleted were reserved at both ends of Neo gene as homologous arms, so as to undergo homologous recombination with genes on chromosome (Figure 2a).

### Identification of mouse genotype

Genomic DNA were extracted from tail tissues according to a method described by Wang^32^, then *Mal* gene was amplified by PCR technology. The PCR products were analyzed by 1.5% agarose electrophoresis, *Mal*+/+, *Mal*+/- and *Mal*-/-mice were identified according to the product size. The primers for *Mal* were as follows: forward primer P1 5’-AACAGGCACTCACTCAGAAGAAAGGGTAG-3’, reverse primer P2 5’-GAGAGCGTGCTTAGATGGTAGTCAC-3’; reverse primer P3 5’-TCGCATTGTCTGAGTAGGTGTCAT-3’. The two primers P2 and P3 were shared the same forward primer P1. The PCR product was 1060 bp of P1 and P2 and 634 bp of P1 and P3(Figure 2b).

### Oral tumor induction

In our study, chemical carcinogen 4NQO dissolved in drinking water was used to induce oral tumor. *Mal*+/+ and *Mal*-/- mice aged more than 8 week were treated with 4NQO at the dose of 40 μg /mL for 20 weeks, then the mice were treated with normal water for another 4 weeks. As for the dose of 80 μg /mL, the mice were fed with 4NQO water for 10 weeks, and then with normal water for another 16 weeks. The mental state and activity of the mice were observed twice a week during the experiment, the color, gloss and presence of oral lesions of mice were observed under strict compliance with the animal protection and welfare conditions. At the same time, the occurrence time, number and growth rate of oral lesion was recorded.

### HE staining

The mice were sacrificed, then tongue tissues, esophagus, spleen, thymus, kidney, liver, lung, skin, stomach and testis were fixed in 4% paraformaldehyde more than 24 hours. Next, the samples were dehydrated, embedded in paraffin and sectioned. The slices were dewaxed, hydrated, stained with hematoxylin and eosin, dehydrated and finally sealed.

### Immunohistochemistry staining

The sections were deparaffinized, rehydrated, submerged into citric acid buffer for heat-induced antigen retrieval, immersed in 0.3% hydrogen peroxide to block endogenous peroxidase activity, blocked with 3% bovine serum albumin. The sections of mice were incubated with anti-Ki67 (#GB111141, Servicebio, 1:500) and anti-Cyclin D1(#GB130719, Servicebio, 1:200) and the sections from patient samples were incubated with antibodies including anti-MRP8 (Abcam, #ab92331,1:200), anti-Aquaporin 3 (Abcam, #ab125219, 1:200), anti-Desmoglein 1 (Abcam, #ab124798, 1:200) and anti-MAL (Santa Cruz, #sc-390687, 1:50) overnight at 4□. The sections were immersed by secondary antibody at room temperature for 1 hour the following day. The sections were then counterstained with hematoxylin, dehydrated, cleared and mounted. The IHC staining of mouse tumor tissue was evaluated by Image J software (v 1.4.3.x). The staining intensity was normalized and measured by the IHC_Profiler plugin integrated into the program. Up to 3 vision fields were selected and calculated to represent the average strength of IHC, the process was repeated on different slides from 3 mice and recorded to examine the statistic significance.

### Biochemical examination of blood

Before sacrificed the mice, venous blood was collected. In one hand, part of the blood was sent to detect blood routine and biochemical indexes such as liver and kidney function. In the other hand, the red blood cell were removed by using reticulocyte lysate and the immune cells were detected by flow cytometry.

### Flow cytometry

The spleen tissues were grinded, and suspended with PBS, then filtered with 70 μm cell strainer. The content of T cells, B cells and NK cells was analyzed using a BD FACS Calibur flow cytometer (BD Biosciences). Conjugated fluorescent antibodies were added to a single-cell suspension. Following incubation with antibody on ice for 30 min in the dark, the cells were washed twice with PBS and analyzed by flow cytometry. The following fluorochrome-conjugated monoclonal antibodies were used according to the manufacturer’s instructions: anti-CD3(#555274, BD Pharmingen), anti-CD4(#553052, BD Pharmingen), anti-CD45R (#561881, BD Biosciences), anti-CD49d (#553157, BD Biosciences), anti-NK1.1 (#108710, Biolegend).

### Tissue dissociation and cell purification

The induced oral tumor tissues were dissected and transported in sterile culture dish with 10 mL 1×Dulbecco’s Phosphate-Buffered Saline (DPBS; Thermo Fisher, Cat. no. 14190144) on ice to remove the residual tissue storage solution, then minced on ice. The dissociation enzyme 0.25% Trypsin (Thermo Fisher, Cat. no. 25200-072) and 10 μg/mL lDNase I (Sigma, Cat. no. 11284932001) dissolved in PBS with 5% Fetal Bovine Serum (FBS; Thermo Fisher, Cat. no. SV30087.02) was used to digest the tissues. Oral tumor tissues were dissociated at 37 □ with a shaking speed of 50 rpm for about 40 min. We repeatedly collected the dissociated cells at interval of 20 min to increase cell yield and viability. Cell suspensions were filtered using a 70 μm nylon cell strainer and red blood cells were removed by 1×Red Blood Cell Lysis Solution (Thermo Fisher, Cat. no. 00-4333-57). Dissociated cells were washed with 1 × DPBS containing 2% FBS. Cells were stained with 0.4% Trypan blue (Thermo Fisher, Cat. no. 14190144) to check the viability on Countess® II Automated Cell Counter (Thermo Fisher)

### Single-cell RNA preparation and sequencing

Beads with unique molecular identifier (UMI) and cell barcodes were loaded close to saturation, so that each cell was paired with a bead in a Gel Beads-in-emulsion(GEM) via the 10× Genomics Chromium machine system. After exposure to cell lysis buffer, polyadenylated RNA molecules hybridized to the beads. The GEM was transferred into PCR for reverse transcription. The GEM system contains free poly(dT) reverse primer, CCC was added at the end of reverse transcription, and the gel magnetic bead contains template conversion Oligo (TSO) primer (including rGrGrG), finally making the RNA in the cell be reversely transcribed into a cDNA chain with Barcode and UMI information. All the remaining procedures including the library construction were performed according to the Illumina User Guide. Sequencing libraries were quantified using a High Sensitivity DNA Chip (Agilent) on a Bioanalyzer 2100 and the Qubit High Sensitivity DNA Assay (Thermo Fisher Scientific). The libraries were sequenced on NovaSeq6000 (Illumina) using 2×150 chemistry.

### Statistical analysis and data visualization

All statistical analyses were performed in R v4.1.0. Student’s *t*-test was used to evaluate the statistic significance of IHC score between wildtype and *Mal* knockout mice. The comparison of gene expression between two different groups was performed using Wilcoxon rank-sum tests. Gene expression associated with OS were analyzed with the Kaplan-Meier methods. The scRNA-seq data was interpreted using Seurat (v 4.0.3), monocle (v 2.20.0) and inferCNV (v 1.8.1). The visualization of the data was performed using the ‘ggplot2’ and ‘ggsci’ packages. All hypothetical tests were two-side, and *P*-values < 0.05 were considered significant in all tests.

## Supplementary materials

1. Supplementary Figures.
2. Supplementary Figure legends.

